# Bioimage postprocessing based on discrete wavelet transform and Lucy-Richardson deconvolution methods

**DOI:** 10.1101/2021.07.14.452302

**Authors:** Haoxin Bai, Bingchen Che, Tianyun Zhao, Wei Zhao, Kaige Wang, Ce Zhang, Jintao Bai

**Author notes:** (Wei Zhao, Ce Zhang).

## Abstract

Accompanied with the increasing requirements of probing micro/nanoscopic structures of biological samples, a variety of image processing algorithms have been developed for visualization or to facilitate data analysis. However, it remains challenging to enhance both the signal-to-noise ratio and image resolution using a single algorithm. In this investigation, we propose an approach utilizing discrete wavelet transform (DWT) in combination with Lucy-Richardson (LR) deconvolution (DWDC). Our results demonstrate that the signal-to-noise ratio and resolution of live cell’s microtubule network are considerably improved, allowing recognition of features as small as 120 nm. Notably, the approach is independent of imaging system and shows robustness in processing fibrous structures, e.g. the cytoskeleton networks.

## I. Introduction

### A. Research Background

Nowadays, facing the explosion of biomedical data, automatic image processing using machine learning and artificial intelligence is of growing importance [1-5]. The development of such vision machine is, however, hindered by the varied image quantities obtained in different microscopic setups.

Many algorithms have been developed to improve the spatial resolution and signal-to-noise ratio (SNR) of biological images, including degenerate-model-based algorithms (e.g. deconvolution [6-11]), mathematical transformation-based algorithms (e.g. spectrum analysis [12, 13], DWT analysis [14-18]), and machine-learning-based algorithms (e.g. deep learning [19, 20]), etc. While, most of these algorithms are capable of fulfilling only a single task, e.g., inhibiting noise, identifying structure contours, or improving resolution, which requires the target images to be clear with minor contribution of noise or aberrations.

Nevertheless, the representative features in biological samples are often small, irregular and influenced by strong noise background. For example, the microtubule of fibroblast forms densely packed network [21]. It is difficult to distinguish a single microtubule filament and track its dynamics during various biological processes. For the isotropic or quasi-isotropic features, e.g., the round-shaped and nanometer-sized exosomes, deconvolution-based algorithm can effectively improve the structural resolution [22, 23]. While, for densely packed networks (e.g., the microtubule), fluorescence signal due to emitted background light and autofluorescence originating from areas above and below the focal plane can decrease the signal-to-noise ratio. To the best of our knowledge, there is currently no effective approach to distinguish filament-like or branch-like structures, and simultaneously achieve super-resolution and high signal-to-noise ratio [24].

### A. Previous Works

Most algorithms are developed based on deconvolution methods, e.g. Lucy-Richardson (LR) algorithm [25-29], the Fast Thresholded Landweber (FTL) algorithm, the Generalized Expectation Maximization (GEM) algorithm, etc. In 2006, Bioucas-Dias et al. advanced the GEM algorithm to process macroscopic image [30]. Although he found that the GEM method could improve image quality, the algorithm only compares the SNR before and after processing, which cannot ensure the original image intensity distribution before and after image processing. FTL algorithm is a fast variational deconvolution algorithm, that minimizes a quadratic data term. Vonesch et al. used FTL to process confocal images of a neuron cell [31, 32]. They found that FTL algorithm could achieve an 8 dB improvement in 10 iterations with an insignificant increase in the image SNR, however, deconvolution methods by themself may lead to over-processing and spurious images.

Wavelet method was primarily applied for denoising, e.g., the Expectation Maximization (EM) algorithm [33, 34]. EM algorithm utilizes both wavelet transform and fast Fourier transform to improve the SNR of the image. It can increase the SNR of macroscopic image from 3 dB to ∼ 7 dB after 8 to 10 iterations. Nevertheless, to the best of our knowledge, wavelet method has never been applied to improve image resolution.

## II. Method Principle And Process

### A. Target of Image Process

In this investigation, we demonstrate that by combining DWT and Lucy-Richardson deconvolution methods (DWDC), the spatial resolution of a typical biological image (high noise, blurred and unclear) can be increased to super-resolution level with improved SNR.

Fig. 1a shows the confocal fluorescence image of 3T3 fibroblasts microtubule networks, which were taken using Nikon A1 microscope and Olympus 100X oil immersion lens (NA 1.4). The excitation light wavelength is 640 nm, and the emission peak is around 674 nm for the SiR-Tubulin dye. Each fluorescence image has 512×512 pixels, with a dot pitch of 0.25 µm. For 3D reconstruction, a total of 20 images were captured by z-stacking, with 1 µm vertical interval. It is obvious that the branch-like microtubule structures are highly contaminated by noise and the structures are clearly bold (Fig. 1b and 1c). Structural features reflecting cell-cell interactions are indistinguishable.

**Fig. 1.**
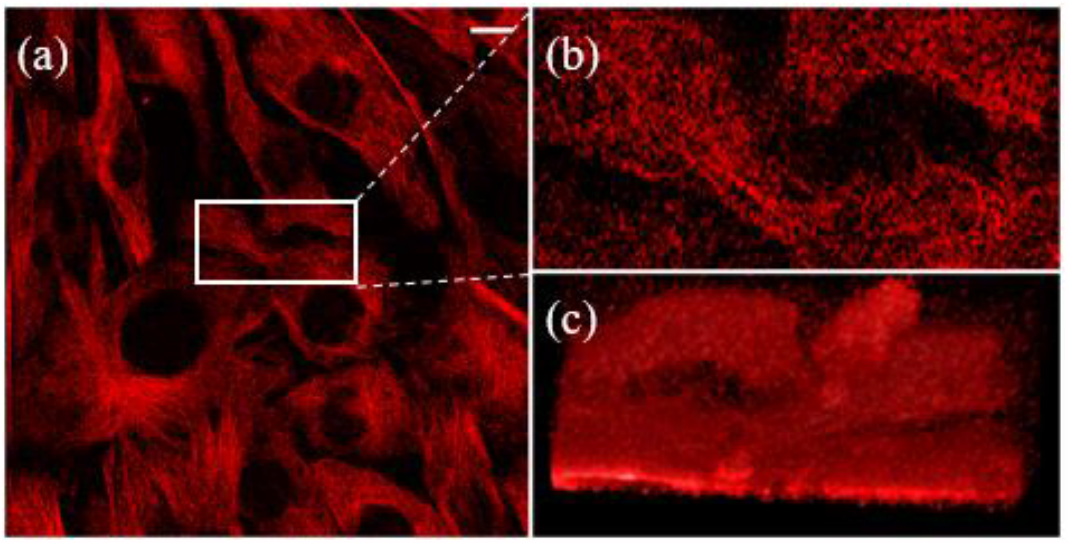
Original confocal image of 3t3 cell microtubule. (a) Image has 512×512 pixels, with a dot pitch of 250 nm. (b) The image in white box. (c) The three-dimensional (3D) reconstruction of (b). The white scale bar represents 10 µm.

### B. Methods and Process

An optical image is a convolution of object with the PSF of the optical system [35]. If *M* is the matrix of the image,

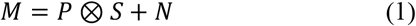

where *P* is the PSF of the optical system, *S* is the light distribution according to the object and *N* is the measurement noise of the optical system. If the size of the PSF is larger than the size of the mesostructure of the actual object, the imaging result has an insufficient spatial resolution to reveal the detail of the original object. Accordingly, the image after the optical system is blurred relative to the actual object.

The DWDC method advanced in this investigation utilizes both LR and DWT, as diagrammed in Fig. 2. Firstly, the image was processed using Gaussian interpolation and threshold analysis. DWT was then applied to suppress noise level and extracts characteristic microtubule structures on the basis of scale analysis. Subsequently, outline of the representative structures is distinguished by binarization with threshold processing, i.e. logical matrix 1. Application of deconvolution method shrinks the outline, and further enhances the spatial resolution. The image is then processed with repeated binarization, threshold analysis and Gaussian interpolation before finalization.

**Fig. 2.**
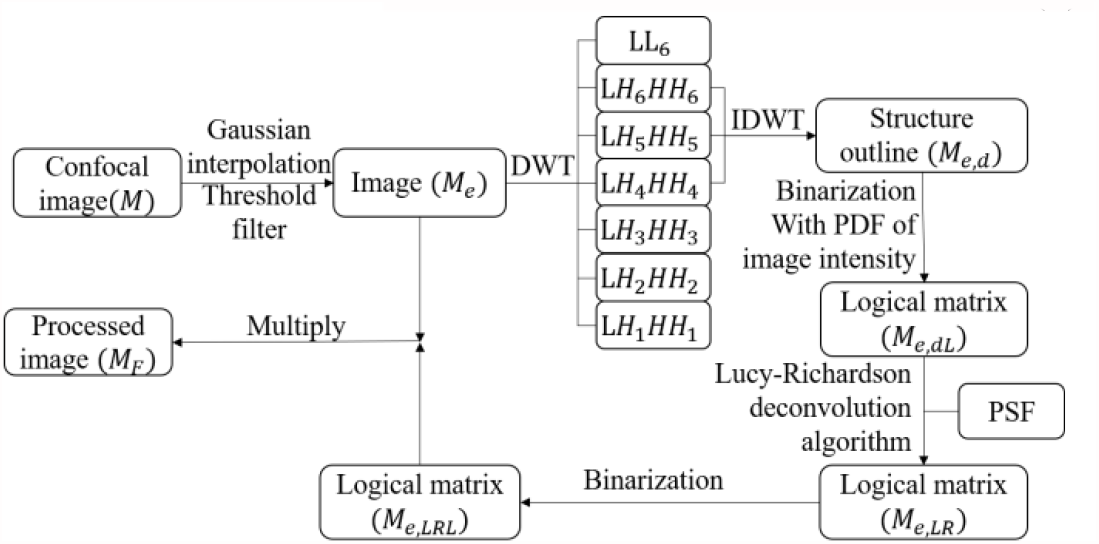
Schematic of DWDC method. Here, we perform Gaussian interpolation and quarter average of image (*M*_*e*_) threshold filtering on the original image. In DWT wavelet processing, *LL*_*n*_ is the approximate wavelet decomposition term, *LH*_*n*_ are the detail wavelet decomposition terms in the x-direction and y-direction, *HH*_*n*_ are the detail wavelet decomposition terms in the diagonal direction. The subscripts of the terms represent the order of wavelet decomposition. During inverse DWT, only 4-6 order terms are included in this investigation. Next, *M*_*e,LRL*_ is obtained from *M*_*e,d*_ after a series of binarization and deconvolution processes. Finally, the overall processed image *M*_*F*_ can be obtained by *M*_*e,LRL*_ ·*M_e,LRL_*.

**Fig. 3.**
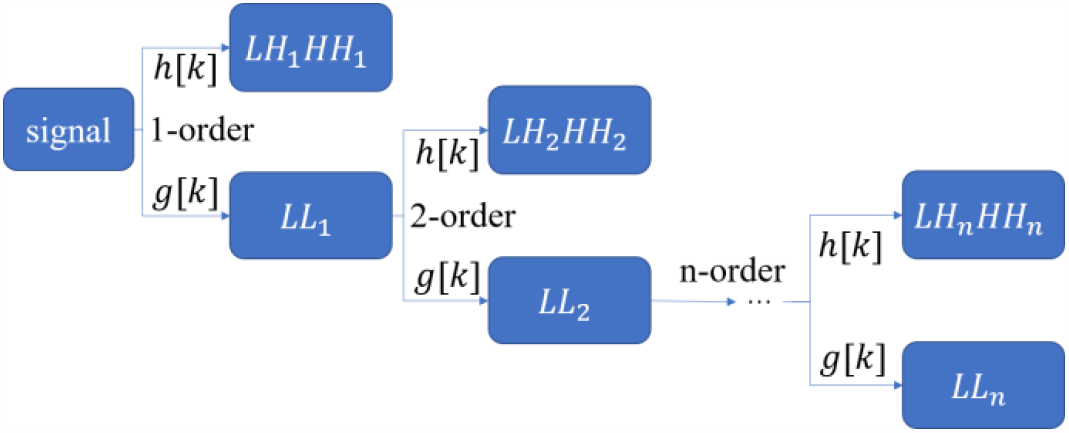
Wavelet decomposition process, where *g*[*k*] is low pass filter, and *h*[*k*] is high pass filter.

#### 1) Expansion of Image by Gaussian Interpolation

To increase resolution of the image, we first reduce the dot pitch. 3D Gaussian interpolation [36] was applied to expand the size of the image with reduced pitch. The original 20 images of 512×512 pixels were expanded twice to 80 expansion images (*M*_*e*_) of 2048×2048 pixels. The Gaussian function is :

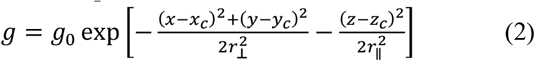

where *r*_⊥_ = (0.61λ_*e*_) /*N_A_* and *r*_∥_ = (4*n*λ_*e*_) / (2*N*_*A*_ ^2^) are the lateral and axial Gaussian radius respectively [37-39], *N*_*A*_ is numerical aperture of lens, λ_*e*_ is the wavelength of the excitation beam. After the expansion, the dot pitch is reduced to 63 nm.

#### 2) Extraction of Microtubule Structures by DWT

The expanded image matrix need to extract microtubule structures by DWT [40], When performing wavelet decomposition, the information corresponding to the scale function is usually filtered by a low-pass filter, and the information corresponding to the wavelet function is filtered by a high-pass filter. At the same time, the scale information obtained by the low-pass filter can be used as the generating function of the wavelet function and the scale function of the next stage. The information corresponding to the scale function represents the low-frequency component in the original signal, which represents the coarse information of the original signal; the information corresponding to the wavelet function represents the high-frequency component in the original signal, which represents the detailed information component of the original signal.

For two-dimensional image matrix, wavelet decomposition will be processed in three directions, i.e. horizontal, vertical and diagonal directions, as:

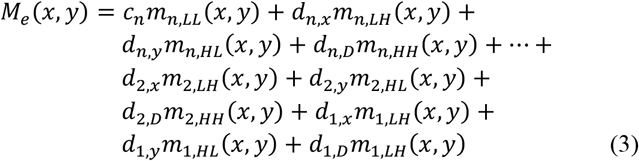

where *c*_*n*_ is the approximate wavelet coefficient after n^th- order discrete wavelet decomposition, *d*_*n,x*_, *d*_*n,y*_ and *d*_*n,D*_ are the detail wavelet coefficients of level n in horizontal, vertical and diagonal directions respectively, as shown in Fig. 4. *m*_*n,LL*_ is the approximate information. *m*_*n,LH*_ is the detailed information in the x-direction. *m*_*n,HL*_ is the detailed information in the y-direction, and *m*_*n,HH*_ is the detail information in the diagonal direction.

**Fig. 4.**
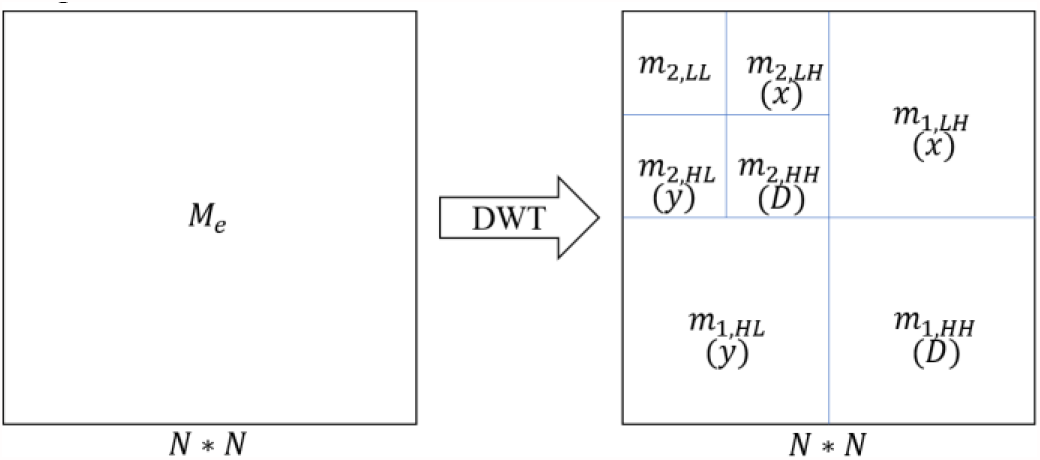
Diagram of second order wavelet decomposition on *M*_*e*_. The left one is the image matrix for decomposition, and the right one shows the decomposed matrices. *N* is the size of the image matrix. *N* = 2048 in this investigation.

They are obtained from the image *M*_*e*_ by the n-order DWT.

*m*_*n,LL*_, *m*_*n,LH*_, *m*_*n,HL*_ and *m*_*n,HH*_ have the formats as follows:

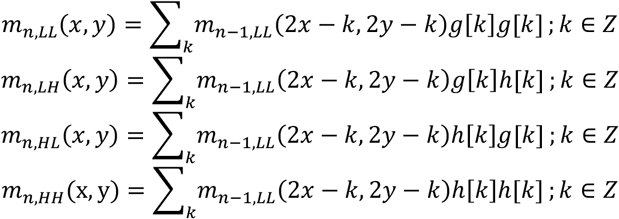

where *m*_0*,LL*_(*x,y*)=*M*_*e*_(*x,y*). After DWT, Eq. 1 can be expressed as:

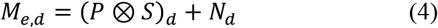

The image information can be further divided into two parts on scale spaces, i.e., the scales related to the demanded structures (denoted as *M*_*e,d*_(*x, y*)) and the scales related to undesired structures (denoted as *M*_*e,ud*_(*x, y*)) which can also be considered as noise structures. Thus,

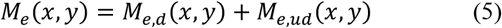

where

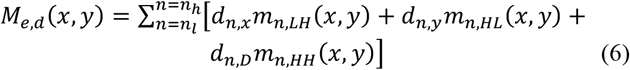

where *n*_*l*_ and *n*_*h*_ correspond to the lower and higher bounds of DWT orders of the demanded structures. For instance, if the demanded structures have a characteristic size between 16 and 64 pixels, we have *n*_*l*_ = 4 and *n*_*h*_ = 6. According to DWT decomposition, the image structures within demanded scale ranges (or frequency ranges) can be retained on each local positions. While the image structures that undesired, e.g. noise (which normally has high-frequency components) and image distortions due to nonuniform illumination (which has low-frequency components), can be removed from the images without the requirement of knowing their detailed distribution.

#### 3) Binarization of Image

When we get the demanded structures extracted by DWT, logic processing was carried out. The threshold value (*χ*) is selected according to the probability density distribution of image intensity. The binarization image is thus obtained as a logical matrix shown below:

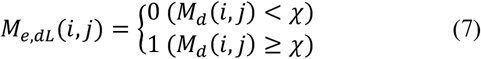

After the processing, the image noise is further inhibited and the outline of demanded structure is highlighted.

#### 4) Resolution Improvement by Lucy-Richardson (LR) Deconvolution Method

The logic matrix obtained in this way is not detailed enough and the structural resolution has not been apparently improved. We then use the Lucy-Richardson deconvolution method to further process the image. Instead of directly applying LR on the image after DWT analysis, in this investigation, we apply LR on the logic matrix (*M*_*e,dL*_) to restore the sketch of the filament structures. This approach can avoid the spreading of high intensity structures that affects the low-intensity structures and leads to spurious images or overprocessing.

LR method is developed on the basis of Bayesian theory [41], Poisson distribution and maximum likelihood estimation. The overall expression formula of LR deconvolution method algorithm is as follows:

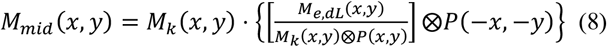

where *M*_*k*_(*x, y*) is the *k*^*th*^ iteration of logic matrix *M*_*e,dL*_ and *M*_*mid*_(*x, y*) is an intermediate result. In each iteration of optimization, a scale factor *f* is applied to evaluate the effect of this processing, according to the image difference before and after the iteration as:

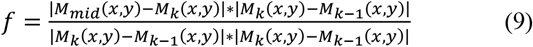

Thus, the *k*^*th*^ iteration of the image can be obtained as:

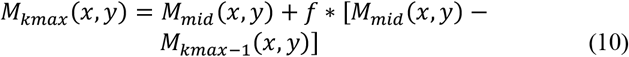

where *kmax* is the maximum number of iterations.

During the deconvolution, one of the most important prerequisites is the evaluation of the actual PSF of the optical system. During LR deconvolution on the image, the PSF of the confocal imaging system is a two-dimensional (2D) Gaussian function, which can be expressed as 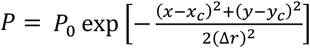, with Δ*r* = ξ*r*_⊥_ being the actual PSF for LR deconvolution and ξ is an experience coefficient which should be determined by a numerical experiment. Furthermore, in the iterative process of deconvolution, background noise can also be gradually amplified, resulting in spurious images or overprocessing. Therefore, optimize the number of iterations (i.e. *kmax*) and threshold deviation (damping coefficient) is important in the application of LR deconvolution. The evaluation of the image after deconvolution is highly arbitrary depends on the demanded structures of the image. In many cases, the restoration quality of an image cannot be simply evaluated by peak signal-to-noise ratio (PSNR) or structural similarity index measure (SSIM). Therefore, in this investigation, the deconvolution parameters are also determined by numerical experiments.

#### 5) Post-Processing

After LR algorithm, the processed logic matrix shows nonlinear characteristics. Thus, we carried out another binarization process on *M*_*kmax*_, with the threshold value *χ* = 1. The secondary logic matrix can be obtained as:

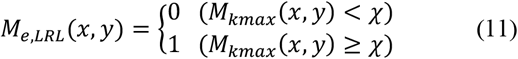

At last, the final result (*M*_*F*_(*x, y*)) of DWDC processing can be obtained by multiplying the expanded image *M*_*e*_ with *M*_*e,LRL*_, to extract the microtubule structure of the 3T3 cell, i.e.)

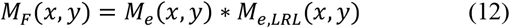

## III. Experimental Results

### A. Expansion of Image by Gaussian Interpolation

A direct comparison between the original 512×512 image (i.e. *M* matrix) and expanded 2048×2048 image (*M*_*e*_) has been carried out in Fig. 5. First, visually, the expansion image shows high consistency as the original one, as shown in Fig. 5(a) and (b). At the corresponding positions, the intensity distributions along the selected row and column both show high similarity (Fig. 5(c, d)), which shows that the details of the original image have been reserved.

**Fig. 5.**
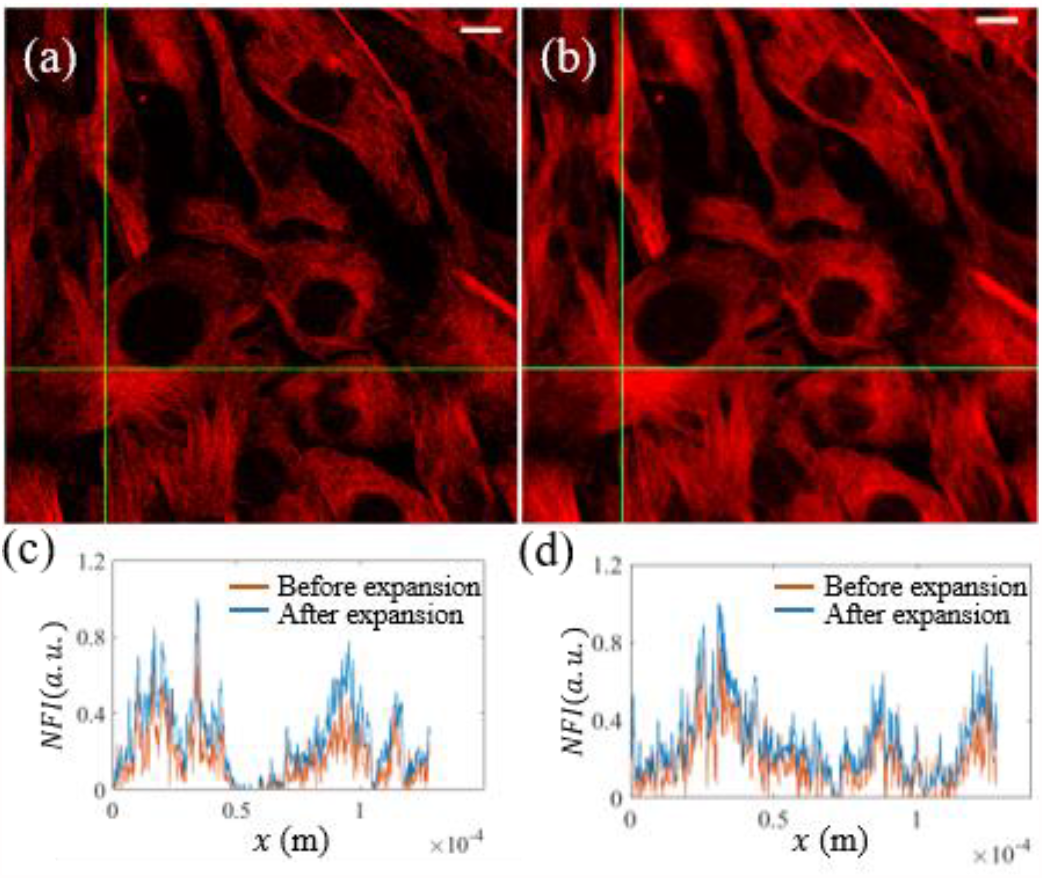
Comparison between the original and expanded images, white scale bars represent 10 µm. (a) Original image; (b) Expanded image; (c -d) Distributions of normalized fluorescent intensity (NFI) on the corresponding horizontal and vertical positions before and after expansion.

### B. Extraction of Microtubule Structures by DWT

In this investigation, we use coif3 wavelet function to decompose the image up to the 6th order. Because the microtubule structure in the image has a width of 20 to 60 pixels, in the extraction, we only keep 4-6 order components, i.e.

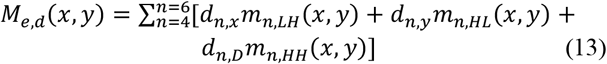

In contrast to the expanded image (Fig. 5(b)), the reconstructed image shown in Fig. 6(a) clearly reserves the filament-like microtubule structure. While the undesired components (Fig. 6(b)), which make the image blur and noisy, have been successfully removed. After DWT analysis, a binarization process was applied according to the probability distribution of *M*_*e,d*_, to extract the sketch of the demand structures. The probability distribution of *M*_*e,d*_ is plotted in Fig. 6(c). Following numerical experiments, we only retain the top 15% of the image intensity. Thus, the corresponding threshold value related to Fig. 6(c) is estimated to be *χ* = 61, where we can clearly see the sketch of the microtubule (Fig. 6(d)).

**Fig. 6.**
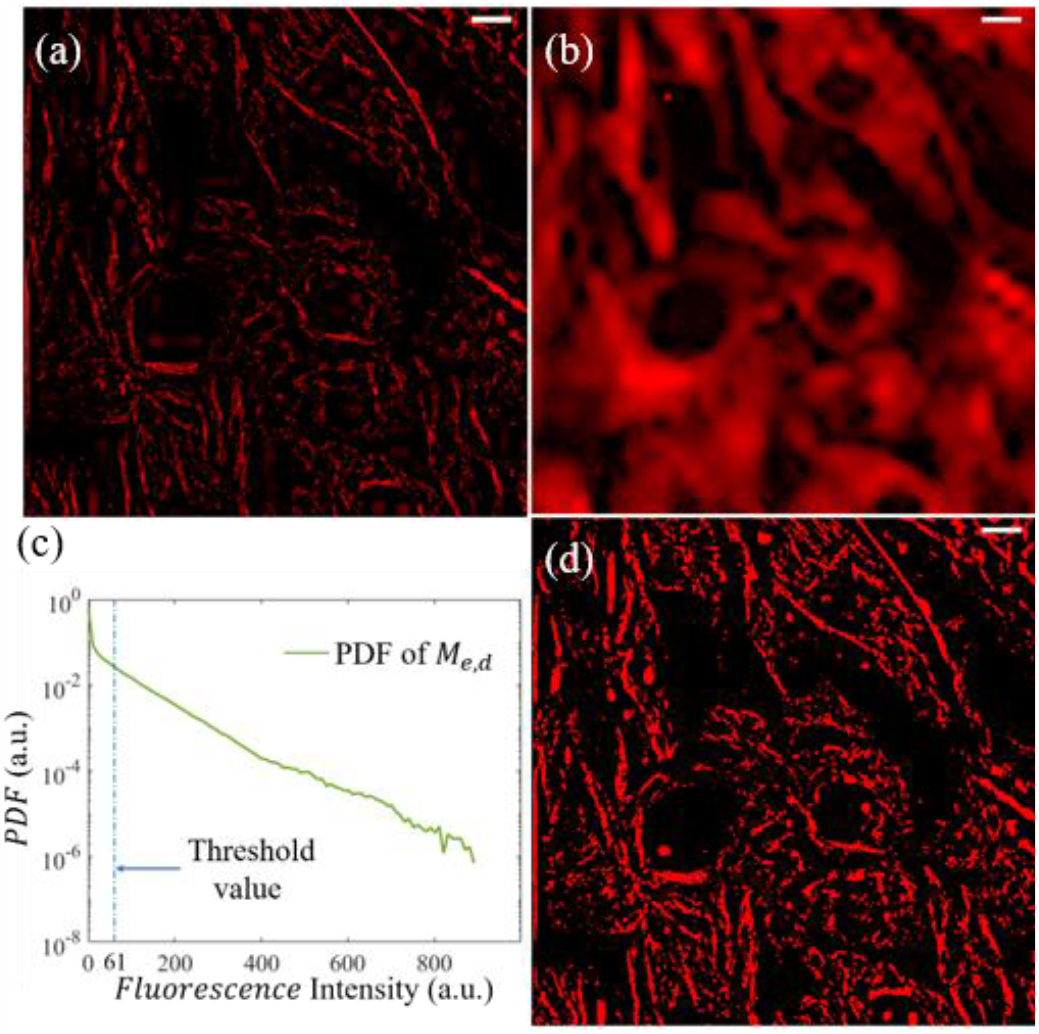
Extraction of microtubule structures by DWT. (a) Extracted components after DWT analysis. (b) Undesired components that should be abandoned. (c) Probability density function (PDF) of *M*_*e,d*_. (d) Logical matrix (*M*_*e,dL*_) of the structure after DWT analysis. The white scale bars represent 10 µm.

### C. Resolution Improvement by LR Deconvolution

At the current stage, the sketch of the structure is still wide, and the spatial resolution of the structures is below our expectation (Fig. 6 (d)), i.e. super resolution. Thereafter, we use LR deconvolution algorithm to further process the logical matrix. During numerical experiments, we tried different ξ, maximum iteration *kmax* and damping coefficient to optimize the outcome (i.e., narrow and consistent structural features). The representative results are listed in Fig. 7(a-d). Notably, all the images in Fig. 7 have been binarized after LR deconvolution.

**Fig. 7.**
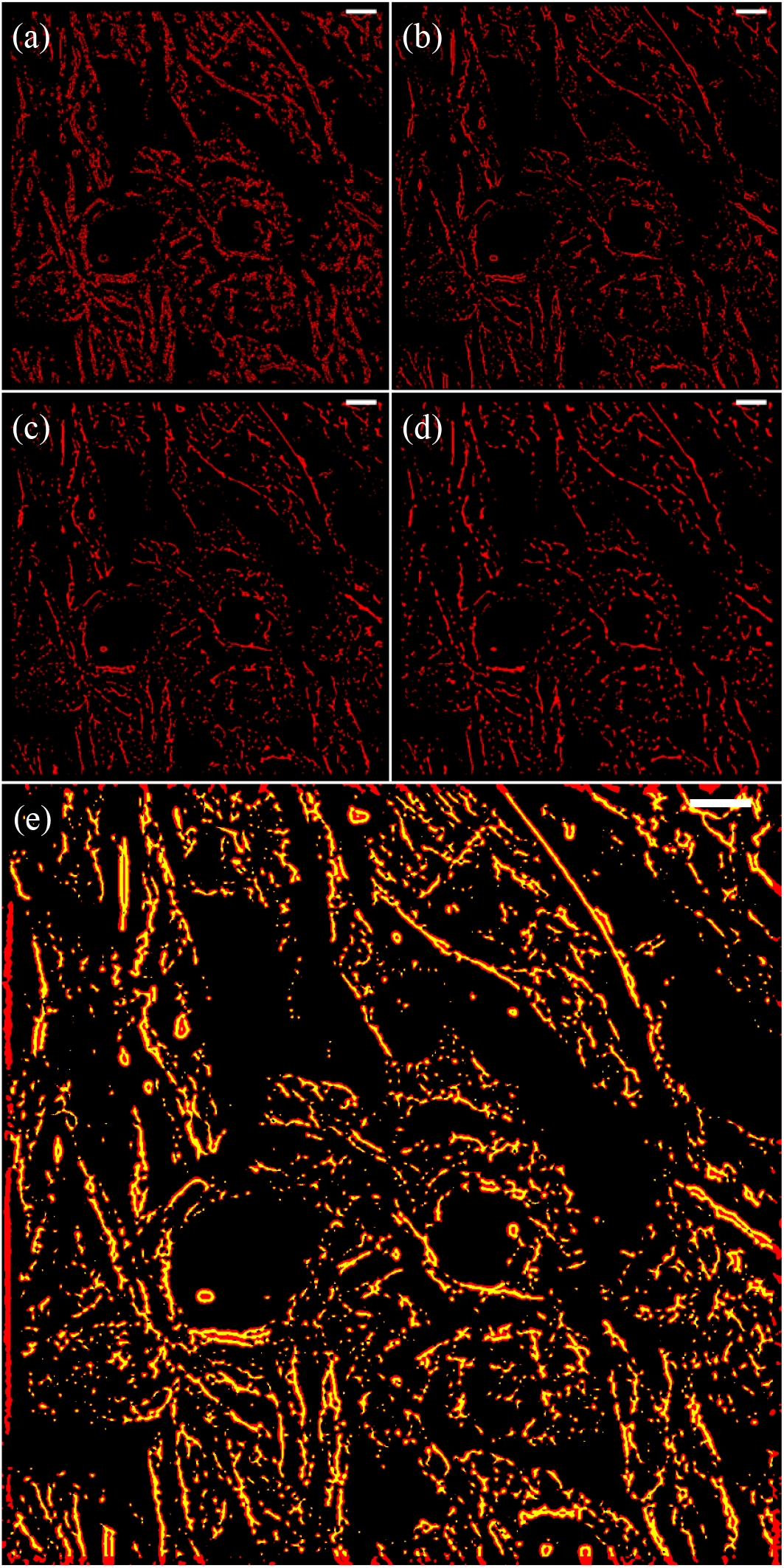
Different empirical coefficients correspond to the deconvolution results, white scale bars represent 10 µm. (a) ξ=0.25; (b) ξ=1.25; (c) ξ=2.25; (d) ξ=3. (e) The comparison between pre-processing and post-processing of the logic matrix used to extract the microtubule structure in the original image. The white scale bars represent 10 µm.

When ξ = 0.5, the applied Gaussian radium for deconvolution Δ*r* is only half of the theoretical value *r*_⊥_. The continuous structures of the logical matrix become fragmented and hollow (Fig. 7(a)). When ξ is increased to 1.25, significant shrinkage of the logical matrix structures was realized with a compromise of continuous network structures (Fig. 7(b)). The optimal outcome was obtained, when *kmax* and damping coefficient are 10 and 0.01 respectively (Fig. 7(e)).

### D. Image After Processing

By extracting the structure of the logical matrix, the microtubule structure of 3T3 cells was obtained (Eq. 12 and Fig. 8(a)). The final image renders the clear filament-like mesh structures of microtubule in 3T3 cells, with ultrahigh contrast and ultralow noise, as shown in Fig. 8(b). A more clear comparison between the original images and processed images can be found from the 3D microtubule structures of 3t3 cells in the supplementary videos.

**Fig. 8.**
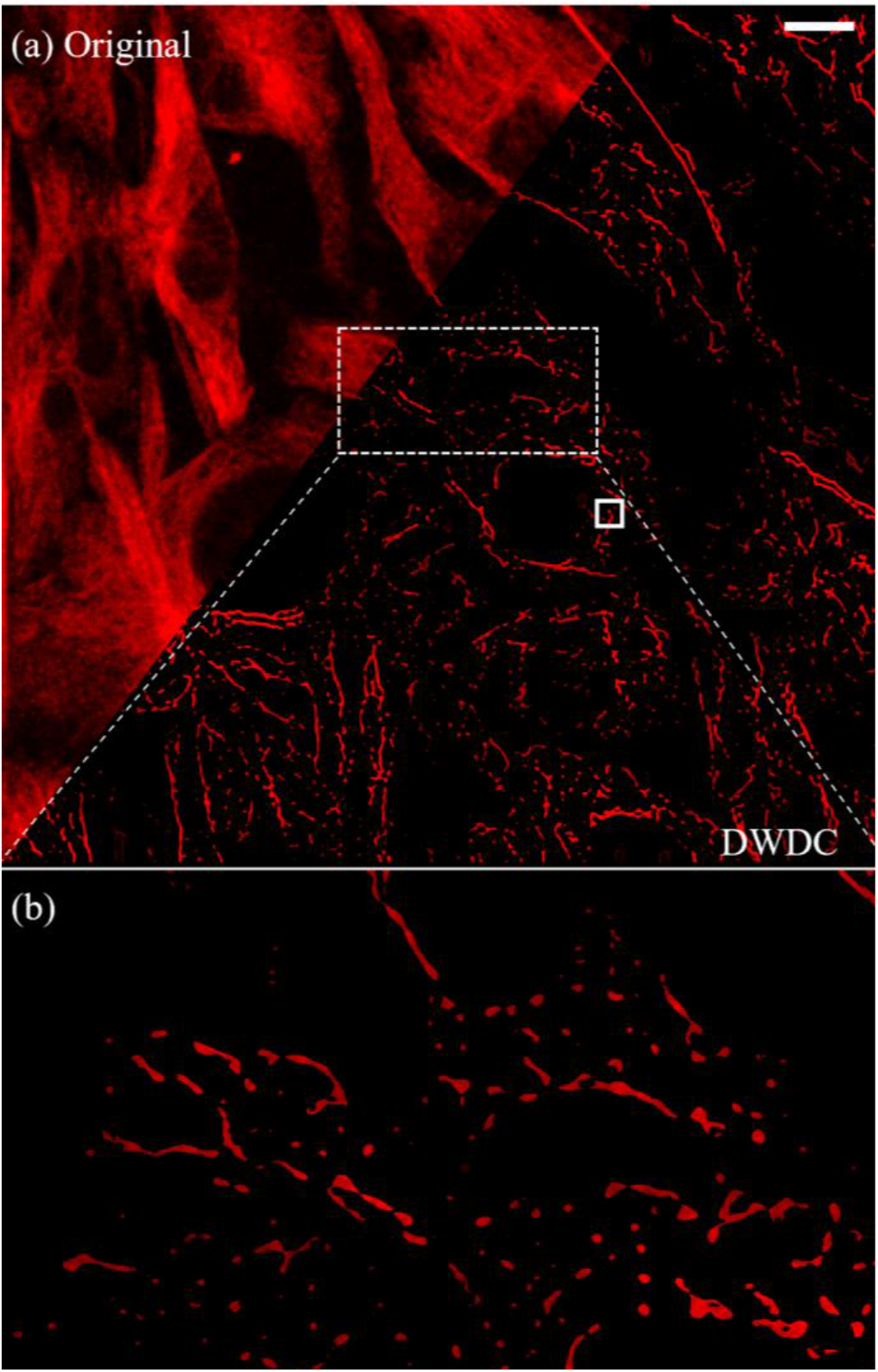
Compare original image and image processed after DWDC method. (a) Direct comparison before and after DWDC method. White scale bar represents 10 µm. (b) Local processing results of DWDC method in the box of dashed line of (a) for comparison with Fig. 1b.

In Fig. 9, we compare the distributions of fluorescence intensity at the same positions of the original and processed images, which shows 15 times improvement in spatial resolution from 1.94 μm to 123.7 nm, as evaluated by the full width at half maximum (FWHM) of the structure.

**Fig. 9.**
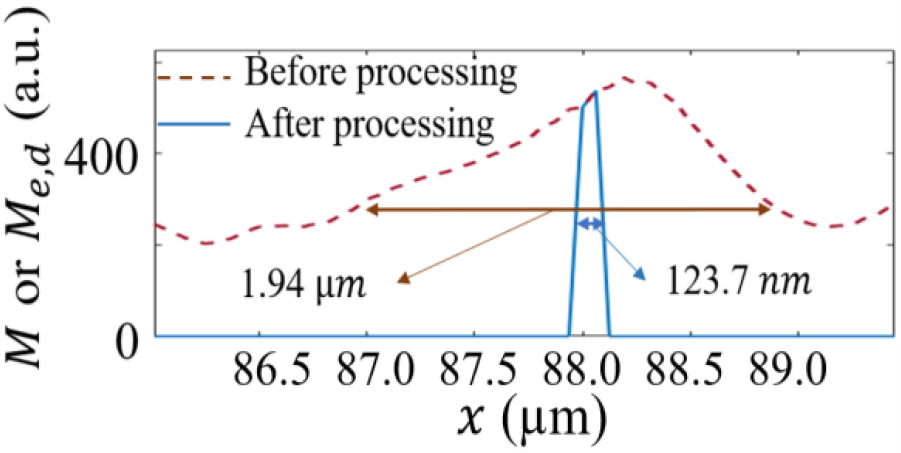
Comparison of FWHM between original image and image processing after DWDC method in the box of solid line in Fig. 8(a). *M* and *M*_*e,d*_ are the image intensities of the 3t3 cell image.

Notably, the PSNR, a commonly used criterion for evaluating the noise level of the images before and after processing is -48.8. SSIM, which is another widely used parameter, is only 0.0155. This implies PSNR and SSIM, as the common judging standards, may not be a golden rule for all the image processing methods.

After processing, the previously unclear images of microtubule network structure (Fig. 1(a) and 1(b)) become considerably more distinguishable (Fig. 10(a)). More biological information has been revealed. As an example, our results demonstrate that 2 cells (i.e., the green- and purple-colored ones) form cell-cell connection. When the third cell (yellow) passes though the gap in between those 2 cells, remodeling of its microtubule network is observed, indicating mechanical forces induced by cell-cell collision [42]. Since it is widely accepted that propagating mechanical cues during collective movement of population cells would activate mechano-signaling and regulate cellular behavior [43], in which remodeling of cytoskeleton networks plays important roles, our approach show potential in deciphering the dynamic cytoskeleton network reorganization and remodeling at the single molecular level, even by a conventional high resolution imaging techniques.

**Fig. 10.**
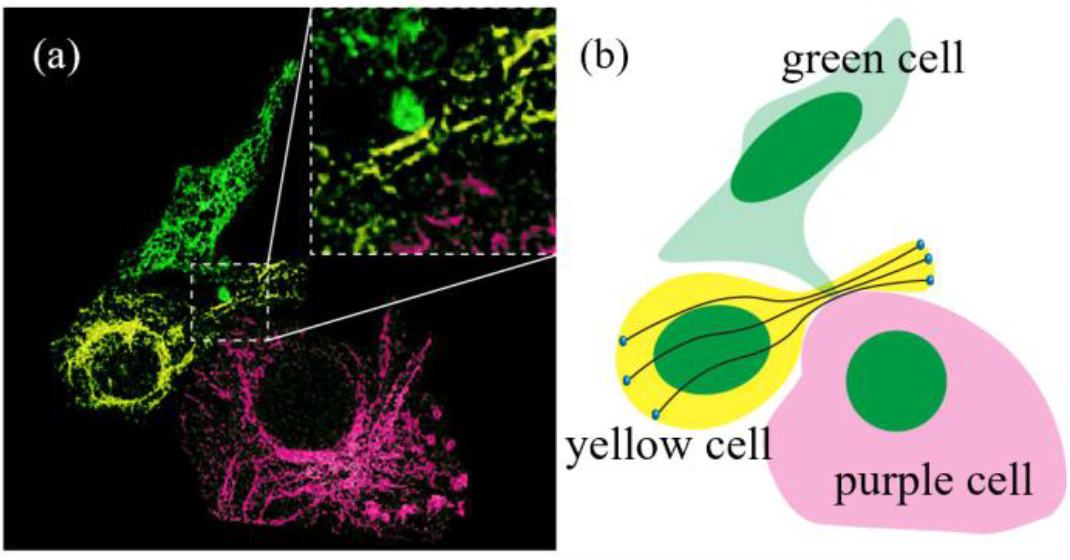
Schematic of cell collision and crossing.

## IV. Conclusions

In this investigation, we introduce DWDC method which is developed based on the discrete wavelet transform and Lucy-Richardson algorithm to extract the microtubule structure of 3t3 cells from confocal images. The microtubule structure in the original image, which has FWHM of up to 1.94 µm, can be reduced to 123.7 nm after processing with DWDC method. The improvement of structural resolution is around 15 times. Compared with the single use of discrete wavelet transform or Lucy-Richardson algorithm for image processing, the composite image processing method can effectively remove noise, improve the SNR and increase the resolution of the image to a super-resolution level simultaneously.

Theoretically, DWDC method is not limited to a certain imaging technique. It identifies the image intensity distribution as long as the scale information of the target structure and the parameter information of the imaging system are known. Thus, this method can become a universal post-processing method for wide-field fluorescence microscope, confocal microscope, STED microscope and etc.

## Supporting information

3D microtubule structure of 3t3 cells by original confocal images

3D microtubule structure of 3t3 cells by DCDW processed images

